# Modeling Combination Chemo-Immunotherapy for Heterogeneous Tumors

**DOI:** 10.1101/2024.01.07.574576

**Authors:** Shaoqing Chen, Zheng Hu, Da Zhou

## Abstract

Hypermutable cancers create opportunities for the development of various immunotherapies, such as immune checkpoint blockade (ICB) therapy. However, emergent studies have revealed that many hypermutated tumors have poor prognosis due to heterogeneous tumor antigen landscapes, yet the underlying mechanisms remain poorly understood. To address this issue, we developed mathematical models to explore the impact of combining chemotherapy and ICB therapy on heterogeneous tumors. Our results uncover how chemotherapy reduces antigenic heterogeneity, creating improved immunological conditions within tumors, which, in turn, enhances the therapeutic effect when combined with ICB. Furthermore, our results show that the recovery of the immune system after chemotherapy is crucial for enhancing the response to chemo-ICB combination therapy.

**Author summary:** The challenge posed by intratumoral heterogeneity is gaining recognition in the field of cancer treatment. Despite the success of immune checkpoint blockade (ICB) therapies in enhancing overall survival across various cancer types, the complexity of therapeutic responses persists due to the heterogeneity of tumor antigens. In this study, we developed mathematical models to explore the evolutionary dynamics of tumors with both homogeneous and heterogeneous antigen landscapes. Our analysis reveals that tumors with heterogeneity exhibit resistance to ICB therapy, unlike their homogeneous counterparts which respond positively. Additionally, our models demonstrate that early treatment of heterogeneous tumors with chemotherapy leads to significant remission but also rapid recurrence. Notably, we identified a fascinating trade-off associated with chemotherapy—while suppressing the immune system, it creates a tumor immunological environment that becomes more conducive to ICB therapy. Finally, our modeling highlights the augmented response observed in tumors subjected to a chemo-ICB combination and shows the crucial role of immune recovery in the context of combination therapy.

## Introduction

Immunogenic tumors can elicit immune responses that suppress tumor growth [1]. Hypermutated cancers produce large amounts of neoantigens and are often considered highly immunogenic and responsive to immunotherapies [2, 3]. Immune checkpoint blockade (ICB) therapy has shown promising results in improving the survival of patients with various types of malignant tumors [4, 5]. Recent evidence suggests that intratumoral heterogeneity has a significant impact on immunotherapy prognosis [6–8]. Immunogenic tumor antigens do not lead to tumor rejection when the fraction of cells bearing them is low [9, 10]. Cancers, characterized by diverse neoantigens, can be unresponsive to ICB.

The loss of immunologic control has been confirmed as one of the emerging hallmarks of cancer [11]. Immune escape in hypermutatble tumors are assisted by checkpoint ligands overexpression to maintain the imbalance between immune surveillance and cancer growth [12]. Check point antibody inhibitors, such as anti-PD-1/PD-L1, are a novel class of inhibitors that function as a tumor suppressing factor via modulation of immune cell-tumor cell interaction [13]. The effect of ICB treatment is dependent on the auto-immunity to eliminate cancers. With a heterogeneous neoantigen landscape, the negative selection imposed on tumor antigens can be dependent on their frequencies [9, 10]. Therefore, intratumoral antigen heterogeneity can lead to compromised immune predation and ICB therapy response [14, 15]. Meanwhile, Chemotherapy faces a similar treatment challenge. The presence of intratumoral heterogeneity and the resulting variability in patients can lead to resistance against chemotherapy. [16, 17].

To tackle the challenge posed by intratumoral heterogeneity, the development of combination protocols involving chemotherapy and immunotherapy holds promise as a strategy for effective cancer treatment [18–20]. Preliminary clinical studies have indicated enhanced effectiveness when immunotherapy is administered alongside chemotherapy [21]. While chemotherapy is traditionally viewed as immunosuppressive, emerging data suggest that it may also harbor immunostimulatory properties [22, 23]. This dual nature presents the potential to create a favorable immunogenic environment within the tumor microenvironment, a feat challenging to achieve solely through immune cell targeting.

Tumor heterogeneity has also fueled investigations into optimal and adaptive therapy through mathematical modeling [16, 24–27]. Nevertheless, further exploration is essential not only to identify the most suitable partners for immune checkpoint blockade (ICB) but also to determine the optimal regimen for combination therapy [18, 19, 28]. Understanding the governing mechanisms of response to combination therapy is crucial for optimizing treatment strategies, which remains incompletely understood. Theoretical models specifically considering neoantigen heterogeneity in chemo-ICB combination therapy are currently very lacking.

In this study, we aim to characterize the evolving dynamics of antigenically heterogeneous tumors under intrinsic immune selection and various therapies. Through mathematical models, we seek to unveil the unique evolutionary patterns of tumors responding to different therapeutic interventions, providing crucial insights for the design of future combination therapies. Existing literature suggests that tumor cell populations suitable for ICB treatment may exhibit resistance to chemotherapy [29, 30]. Immune-cold tumors typically show limited response to ICB monotherapy [31], but these tumors often display heightened expression of receptors promoting vasculogenesis and angiogenesis, such as vascular endothelial growth factor (VEGF) [32]. To transform these immune-cold tumors into inflamed ones, combination therapy involving diverse treatment modalities, including chemotherapy, emerges as a promising therapeutic strategy. These approaches are actively under investigation [18, 32]. Our focus is on uncovering the distinct mechanisms underlying these dynamics and the intratumoral antigenic profile of tumors. We developed an immunogenic heterogeneity model (IHeM) to explore the impact of combination therapy on heterogeneous tumors. For comparison, we also established a typical immunogenic homogeneity model (IHoM) for antigenically homogeneous tumors. We observed that heterogeneous tumors demonstrated resistance to ICB therapy but exhibited early remission when treated with chemotherapy alone. Notably, the combination of chemotherapy and ICB demonstrated a curative effect on heterogeneous cancers modeled by the IHeM. Our results also revealed that chemotherapy effectively mitigates antigen heterogeneity, enhancing the immunological conditions within cancers and thereby improving the efficacy of combination therapy. Furthermore, our findings highlighted that the recovery of the immune system significantly enhances the response to combination therapy.

## Methods

### Modeling cancer evolution under immune system selection

We consider two distinct models: the immunogenic homogeneity model (IHoM) and the immunogenic heterogeneity model (IHeM). These models are designed to capture the evolutionary dynamics of antigenically homogeneous and heterogeneous tumors, respectively. In order to represent the interactions between cancer cells and the immune system, we employ the predator-prey framework, as described in [33]. In this framework, cancer cells are akin to prey, susceptible to being targeted or attacked by immune cells, which act as predators in the biological system.

In the immunogenic homogeneity model (IHoM), it is assumed that immunogenic cells originate from antigenically neutral cells through the accumulation of antigens. Our model involves a cell population organized into three compartments, each representing a distinct cell type. As illustrated in Fig. 1a, compartment NCs represents antigenically neutral cells, compartment ICs represents immunogenic cells, and compartment ECs represents immune-escaped cells. For NCs, we assume a division rate of *r*_1_. Upon division, a cell can either undergo transformation into one immunogenic cell through the accumulation of antigenic mutations at a rate *μ*_1_ or produce two identical offspring at a rate of 1 − *μ*_1_. Immunogenic cells divide at a rate of *r*_2_ and face a death rate of *d* due to immune system activity. Through the overexpression of immune checkpoint ligands, cancer cells can evade immune elimination of immunogenic cells. Additionally, to account for Immune Checkpoint Blockade (ICB) therapy, we introduce immune escape [9, 34], wherein immunogenic cells can transform into immune-escaped cells (ECs) at a rate of *p*_*e*_ (Fig. 1a). The IHoM delineates the fundamental mechanisms governing tumor evolution under immune system selection. The cellular processes in the IHoM are summarized as follows:

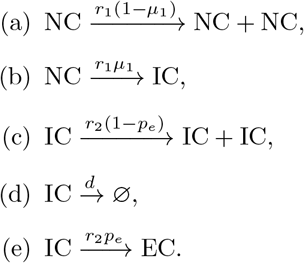

**Fig 1.**
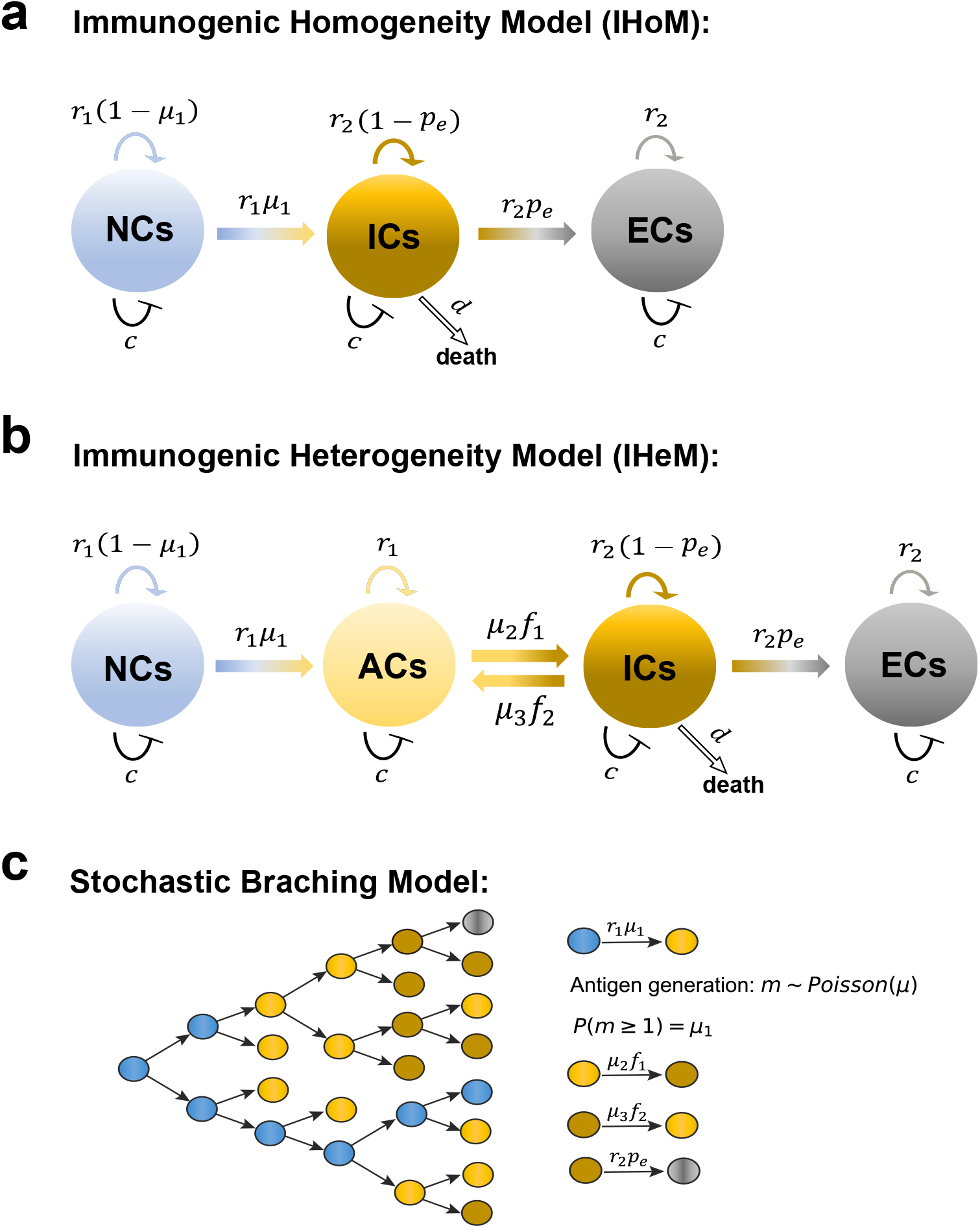
Representation of our models. **a**, Schematic representation of the immunogenic homogeneity model (IHoM). Different compartments represent different cell immunogenic status. From left to right, it represents neutral cells (NCs), immunogenic cells (ICs) and immune escaped cells (ECs). For each NC, it can divide at rate *r*_1_, and then either give birth to two identical offsprings with birth rate 1 − *μ*_1_ or transform into one immunogenic cell at rate *μ*_1_. For each IC, it can divide at rate *r*_2_, and then give birth to two identical offsprings at rate 1 − *p*_*e*_ or transform into an EC at rate *p*_*e*_. ICs die at rate *d*. **b**, Schematic representation of the immunogenic heterogeneity model (IHeM). From left to right, it represents neutral cells (NCs), antigenic cells (ACs), immunogenic cells (ICs) and immune escaped cells (ECs). An AC transforms into an immnogenic cell at rate *μ*_2_*f*_1_. ICs transform into ACs at rate *μ*_3_*f*_2_. Other cellular processes are similar to the ones of the IHoM represented in **a. c**, Schematic presentation of the stochastic branching model. The blue filled circles represent NCs, yellow filled circles represent ACs, blown filled circles represent ICs and grey filled circles represent ECs.

In the immunogenic heterogeneity model (IHeM), we integrate negative frequency-dependent selection into the IHoM [10, 35], as depicted in Fig. 1b. Cells in compartment NCs can initially transform into antigenic cells (ACs) through antigen accumulation. Antigenic cells can proliferate at a rate of *r*_1_ and remain inconspicuous to the immune system when present in low frequencies [9, 10]. A transition from antigenic cells to immunogenic cells (ICs) occurs with a rate of *μ*_2_*f*_1_, where *f*_1_ denotes the frequency of ACs, i.e., the ratio of the number of ACs to the sum of NCs and ACs. To model the impact of tumor antigen heterogeneity on immune recognition ineffectiveness, we assume that immunogenic cells can revert to antigenic cells at a rate of *μ*_3_*f*_2_, where *f*_2_ represents the ratio of the number of ICs to the sum of ICs, NCs and ACs. Neutral and immunogenic cells in the IHeM proliferate and undergo cell death similarly to the IHoM. The cellular processes in the IHeM are summarized as follows:

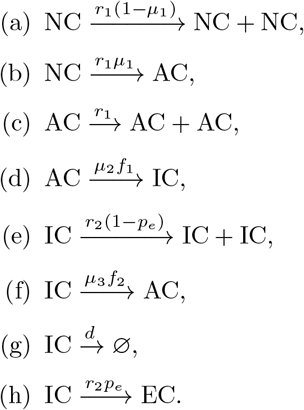

Moreover, competitive interactions between different cells are also considered in our model [36, 37]. We assume that cell competition causes one of the two competing cells to die at equal chance. For example, 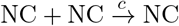 describes the competition between two NCs, and 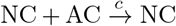 describes the competition between NC and AC. Here *c* is the competition death rate that characterizes the competition strength.

Based on the above assumptions, we can derive the ordinary differential equations (ODEs) of the IHoM and IHeM respectively. For the IHoM,

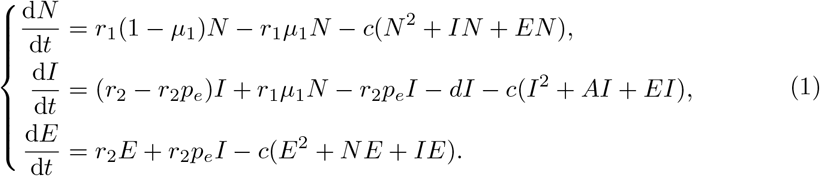

Here, *N, I*, and *E* denote the quantities of NCs, ICs, and ECs, respectively. Neoantigen generation is linked to cell division and is expressed as *r*_1_*μ*_1_*N*. The processes of cell death and immune escape in ICs are represented by *dI* and *r*_2_*p*_*e*_*I*, respectively. Cell competition processes are delineated by terms such as *c*(*N* ^2^ + *IN* + *EN*) at the end of each equation.

For the IHeM,

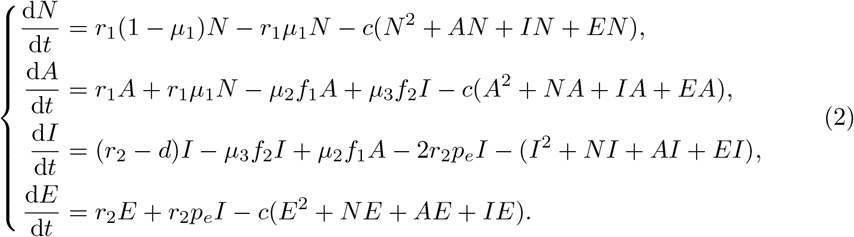

Here, *N, A, I*, and *E* denote the quantities of NCs, ACs, ICs, and ECs, respectively. Neoantigen generation is linked to cell division and is expressed as *r*_1_*μ*_1_*N*. The transitions between ACs and ICs, under negative frequency-dependent selection, are characterized by *μ*_2_*f*_1_*A* and *μ*_3_*f*_2_*I* in the second and third equations, respectively. The processes of cell death and immune escape in ICs are represented by *dI* and *r*_2_*p*_*e*_*I*, respectively. Cell competition processes are delineated by terms such as *c*(*N* ^2^ + *AN* + *IN* + *EN*) at the end of each equation.

To simulate the impact of intratumoral heterogeneity on cancer treatment in greater detail and validate the ODEs model, we conducted stochastic simulations employing a stochastic branching process to depict antigen accumulation during cancer progression (Fig. 1c). In each simulation step, cells that are antigenically neutral and belong to a lineage can proliferate at a rate of *r*_1_. The two daughter cells resulting from each division accumulate antigenic mutations at an overall rate of *μ*. The number of generated mutations, denoted as *m*, is sampled from a Poisson distribution (Poisson(*μ*)), and each antigen’s antigenicity is drawn from an Exponential distribution [34]. In the IHoM, when a cell acquires an antigenic mutation, it transforms into an immunogenic cell, while in the IHeM, it becomes an antigenic cell. Antigenic cells can proliferate at a rate of *r*_1_, and immunogenic cells can proliferate at a rate of *r*_2_. To ensure parameter consistency between the ODEs and stochastic simulations, we select parameters such that *P* (*m* ≥ 1) = *μ*_1_, where *P* (*m* ≥ 1) represents the probability of cells acquiring no less than one neoantigen at cell division (*m* ∼ Poisson(*μ*)), thus we have 1 − *e*^*−μ*^ = *μ*_1_ and then *μ* = − ln(1 − *μ*_1_). Immunogenic cells have a death rate *d* and an immune escape probability *p*_*e*_. All cellular processes align with the ODE models depicted in Fig. 1 a and b. Identical parameter sets are used in both simulations and the ODE models. Furthermore, we introduce cell competition in our simulations to regulate population sizes.

### Modeling ICB therapy and chemotherapy

We model the impact of therapies and the underlying mechanisms governing therapy response based on our IHoM and IHeMs. In various cancer types, Immune Checkpoint Blockade (ICB) therapy has shown enhanced survival for a subset of patients. The effect of ICB is represented in our model by reverting immune-escaped cells back to immunogenic cells (Fig. 2a). To model therapies, we employ impulsive differential equations [33, 38] as follows:

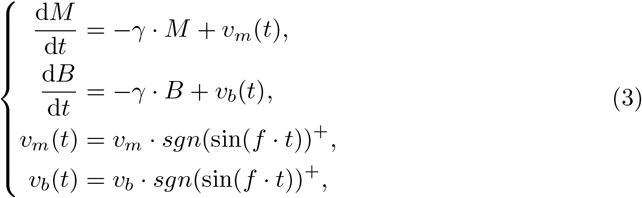

where *M* and *B* represent the blood concentration of chemotherapy and ICB therapy drugs, respectively. *γ* is the drug decay rate, *v*_*m*_ and *v*_*b*_ are the dose of every pulse of drug delivery. In addition,

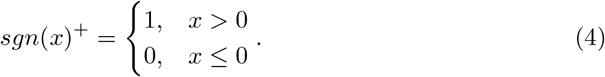

**Fig 2.**
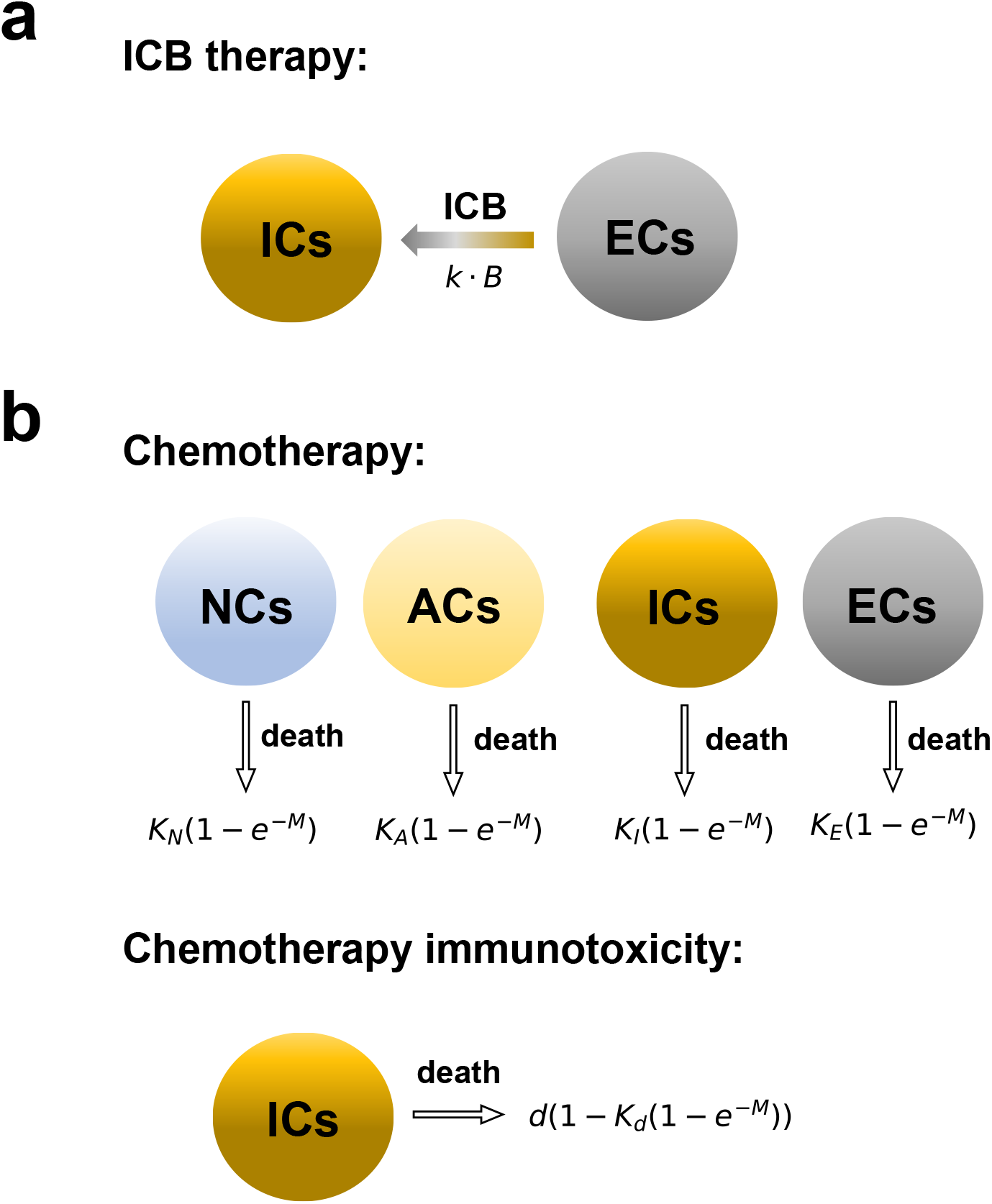
Representation of ICB and chemotherapy models. **a**, Schematic representation of ICB therapy. Here *B* represents the blood concentration of ICB drugs. ICs represent immunogenic cells and ECs represent immune escaped cells. **b**, Schematic representation of chemotherapy. Here *M* represents the blood concentration of chemotherapy drugs. NCs represent neutral cells, ICs represent immunogenic cells, and ECs represent immune escaped cells. The parameters *K*_*N*_, *K*_*A*_, *K*_*I*_, and *K*_*E*_ denote the sensitivity of each cell type to chemotherapy. For example, the death rate of neutral cancer cells under chemotherapy is expressed as *K*_*N*_ (1 − *e*^*−M*^). The parameter *K*_*d*_ represents chemotherapy-induced damage to the immune system (Chemotherapy immunotoxicity), modeled by adjusting the death rate as *d* → *d*. (1 − *K*_*d*_(1 − *e*^*−M*^)).

The parameter *f* governs the therapy cycle, and we define *T* as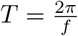, representing the duration of each therapy cycle. During each Immune Checkpoint Blockade (ICB) therapy pulse, a portion (*k*. *B*) of ECs transforms into ICs based on the drug concentration *B*. Consequently, the ODEs describing ICB therapy for the IHoM are given as follows:

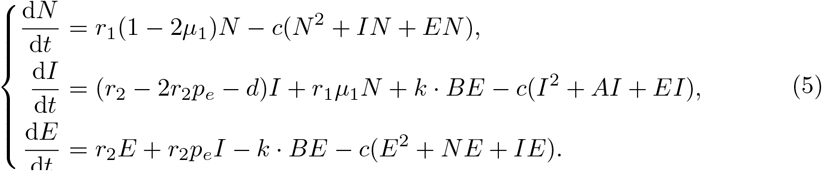

The ODEs for the ICB therapy in the IHeM are provided below:

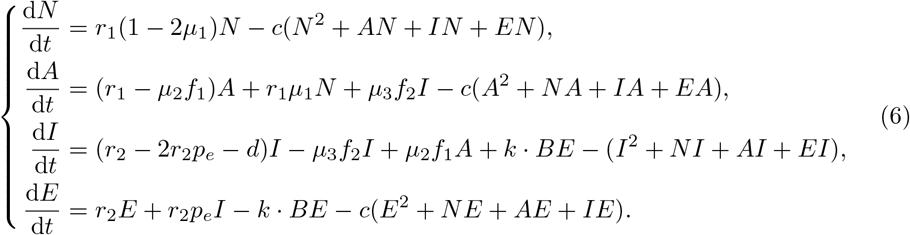

We now incorporate chemotherapy into our model (Fig. 2b). Acknowledging that faster proliferating cancer cells are generally more susceptible to chemotherapy [39, 40], we assume that Neutral Cells (NCs) and Antigenic Cells (ACs) exhibit greater sensitivity to chemotherapy compared to Immunogenic Cells (ICs) and Escape Cells (ECs). We introduce coefficients *K*_*N*_, *K*_*A*_, *K*_*I*_, and *K*_*E*_ to represent each cell type’s sensitivity to chemotherapy. Under chemotherapy, the death rate of neutral cancer cells is given by *r*_1_*K*_*N*_ (1 − *e*^*−M*^) [41]. To align with findings suggesting that cancer cells overexpressing checkpoint ligands are less responsive to chemotherapy [29, 30], we set *K*_*N*_ = *K*_*A*_ = *K*_*I*_ *> K*_*E*_. Considering the known immunosuppressive effects of chemotherapies [33, 42, 43], we also incorporate immunotoxicity of chemotherapy into our models [44]. Let *K*_*d*_ be the coefficient associated with chemotherapy-induced damage to the immune system. We model this damage by adjusting the death rate as *d* → *d* (1 − *K*_*d*_(1 − *e*^*−M*^)).

Consequently, the ODEs describing chemotherapy for the IHoM are:

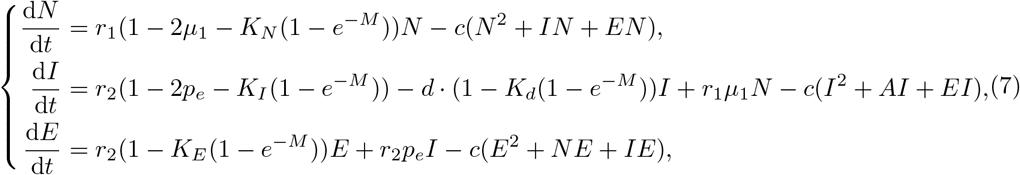

For the IHeM, we have

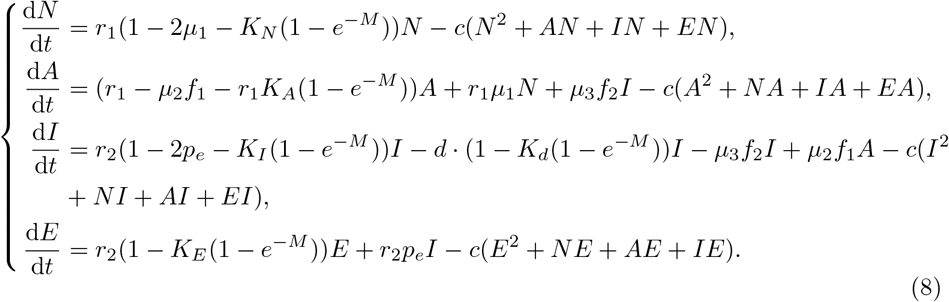

Combining the ICB and chemotherapy models mentioned above, we can establish a model for chemo-ICB combination therapy as follows

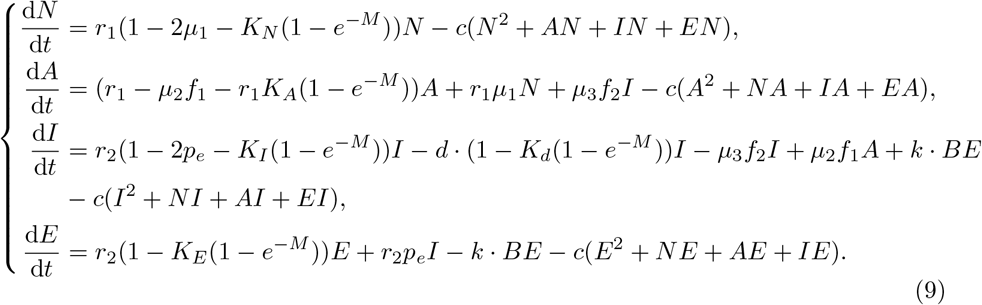

## Results

We initially investigate the growth dynamics of IHeM by employing both ODEs and stochastic simulations. The corresponding parameter values are detailed in Table 1. To study cancer growth in the absence of immune escape, we set *p*_*e*_ = 0. Cancer progression attains equilibrium due to cell competition, as depicted in Figure 3a and b. To assess how the intrinsic strength of immune predation influences cancer, we subsequently examine cancer growth dynamics under the influence of the death rate *d* for immunogenic cells. Our findings indicate that an increase in *d* leads to a decrease in the proportion of immunogenic cells (ICs) and an increase in the proportion of antigenic cells (ACs), as illustrated in Figure 3c. Similar results are observed through stochastic simulations, as shown in Figure 3d and e. The intriguing dynamics observed at equilibrium, highlighting the proportions of antigenic and immunogenic cells, unveil how cancers adapt to auto-immunity within the IHeM. This outcome also offers a plausible explanation for the immune system’s failure to eliminate heterogeneous cancers burdened with a significant antigen load [6, 7, 10].

**Table 1.** Estimated parameter values.

| Parameters | Units | Description | Estimated value |
| --- | --- | --- | --- |
| $r_1$ | day<sup>-1</sup> | Proliferation rate (NCs and ACs) | 1 [34] |
| $r_2$ | day<sup>-1</sup> | Proliferation rate (ICs and ECs) | 0.5 |
| $\mu_1$ | day<sup>-1</sup> | Antigenic mutation rate | 0.5 [4, 6] |
| $\mu_2$ | day<sup>-1</sup> | Immune recognition coefficient | 0.6 |
| $\mu_3$ | day<sup>-1</sup> | Immune exhaustion coefficient | 0.05 |
| $d$ | day<sup>-1</sup> | Cell death rate | [0, 1] |
| $c$ | cell<sup>-1</sup> | Cell competition death rate | $1 \times 10^{-8}$ |
| $p_e$ | day<sup>-1</sup> | Immune escape rate | $1 \times 10^{-5}$ [4, 34] |
| $f$ | day<sup>-1</sup> | Treatment frequency coefficient | $2\pi/21$ [16] |
| $\gamma$ | day<sup>-1</sup> | Rate of drug decay | 0.9 [33, 58] |
| $K_N, K_A, K_I$ | day<sup>-1</sup> | Fractional cell kill by chemotherapy | 0.9 [33, 59] |
| $K_d, K_E$ | day<sup>-1</sup> | Fractional immune cell kill by chemotherapy | 0.6 [33, 59] |
| $v_m$ | pulse<sup>-1</sup> | Dose strength of chemotherapy drug | [1, 5] [33] |
| $v_b$ | pulse<sup>-1</sup> | Dose strength of ICB therapy drug | [0.1, 0.9] |

**Fig 3.**
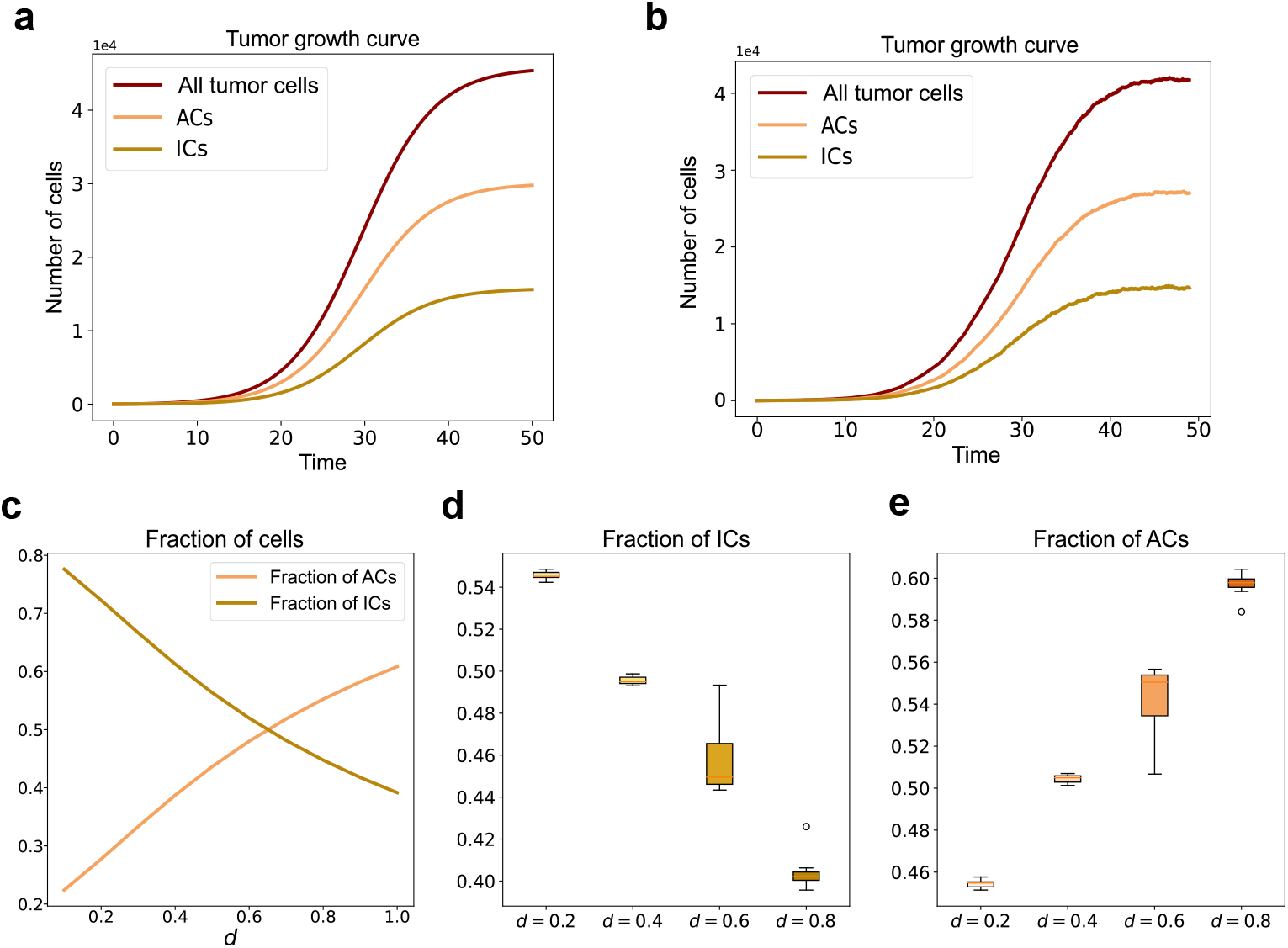
Growth dynamics of the IHeM. **a**, Tumor growth curves generated with numerical solutions of ODEs of the IHeM. Growth curves of ACs, ICs and all tumor cells are shown. **b**, Tumor growth curves generated with simulation data of ACs, ICs and all tumor cells in the IHeM. **c**, Fractions of ACs and ICs generated with numerical solution of ODEs of the IHeM. **d**, Box plots of fractions of ACs generated with 10 simulations of the stochastic IHeM. **e**, Box plots of fractions of ICs generated with 10 simulations of the stochastic IHeM. Parameters: (*r*_1_, *r*_2_, *μ*_1_, *μ*_2_, *μ*_3_, *d, c, p*_*e*_) = (1, 0.5, 0.5, 0.6, 0.05, 1, 1 × 10^*−*5^, 0).

To account for cancer treatment, we introduced immune escape into our models (*p*_*e*_ *>* 0). Our findings reveal that tumors undergo population shifts toward immune-escaped cells, ultimately reaching equilibrium (Fig. 4). Notably, immune escape occurs early in tumors under the IHoM and later in those under the IHeM (Fig. 4). The tumors in these two models follow distinct evolutionary paths before achieving complete immune escape at the whole tumor level. In the IHoM, tumors are swiftly dominated by immune-escaped cells derived from immunogenic cells (ICs), establishing immune escape as the primary mechanism promoting tumor survival. Conversely, in the IHeM, tumors exhibit a capacity for immune selection evasion through high heterogeneity. The significant immune escape observed in hypermutable tumors is considered vital for achieving tumor homogeneity and eliciting an effective immunotherapy response [15, 35]. Nevertheless, our results do not indicate a clear distinction between the IHoM and the IHeM based on this criterion.

**Fig 4.**
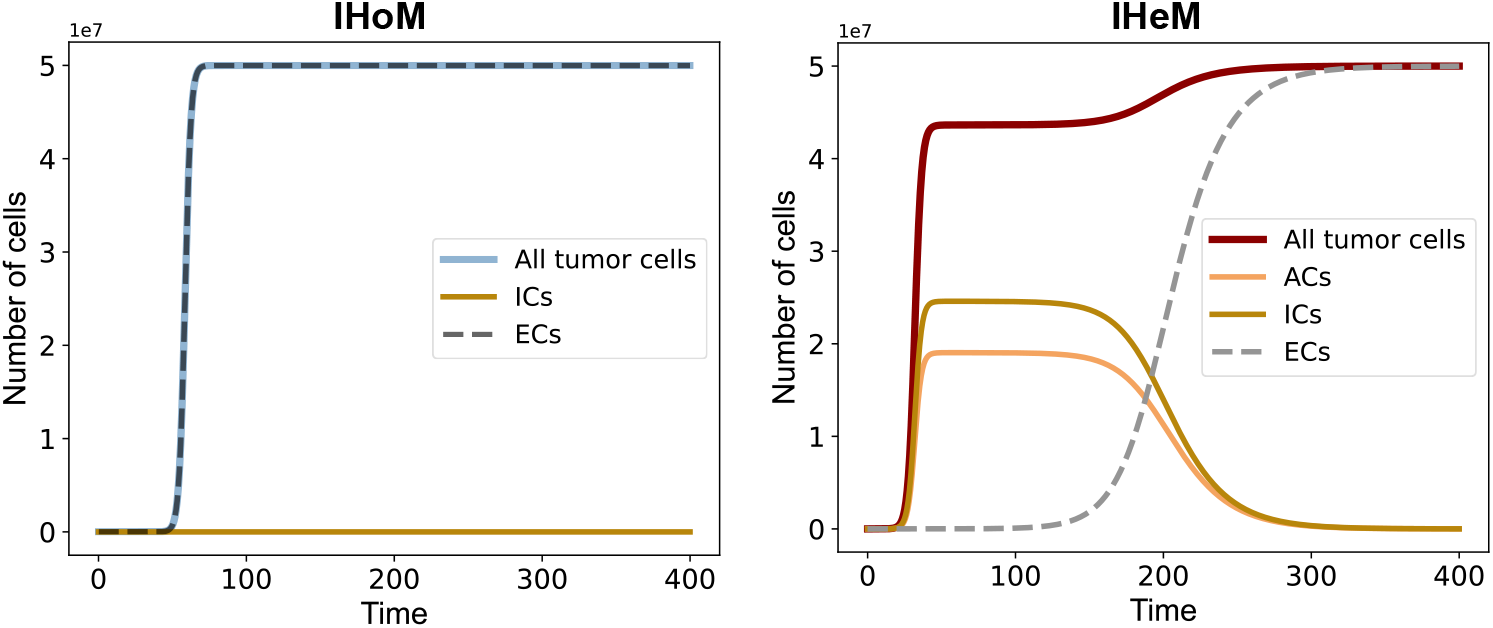
Comparison of the IHoM and IHeM. **a**, Growth curves generated from the numerical solutions of ODEs of the IHoM. **b**, Growth curves generated from the numerical solutions of ODEs of the IHeM. Growth curves of all types of cells are shown. Parameters: (*r*_1_, *r*_2_, *μ*_1_, *μ*_2_, *μ*_3_, *d, c, p*_*e*_) = (1, 0.5, 0.5, 0.6, 0.05, 1, 1 × 10^*−*8^, 1 × 10^*−*5^).

We investigated the impact of immune checkpoint blockade (ICB) and chemotherapy, utilizing standard chemotherapy protocols with a 21-day treatment cycle. Following the conventional practice where drugs are administered during the first half of the cycle, allowing the latter half for patient recovery [16], both chemotherapy and ICB were administered on the same day [45]. Each treatment cycle consisted of 21 days, with one pulse of drugs administered (Fig. 5a and c).

**Fig 5.**
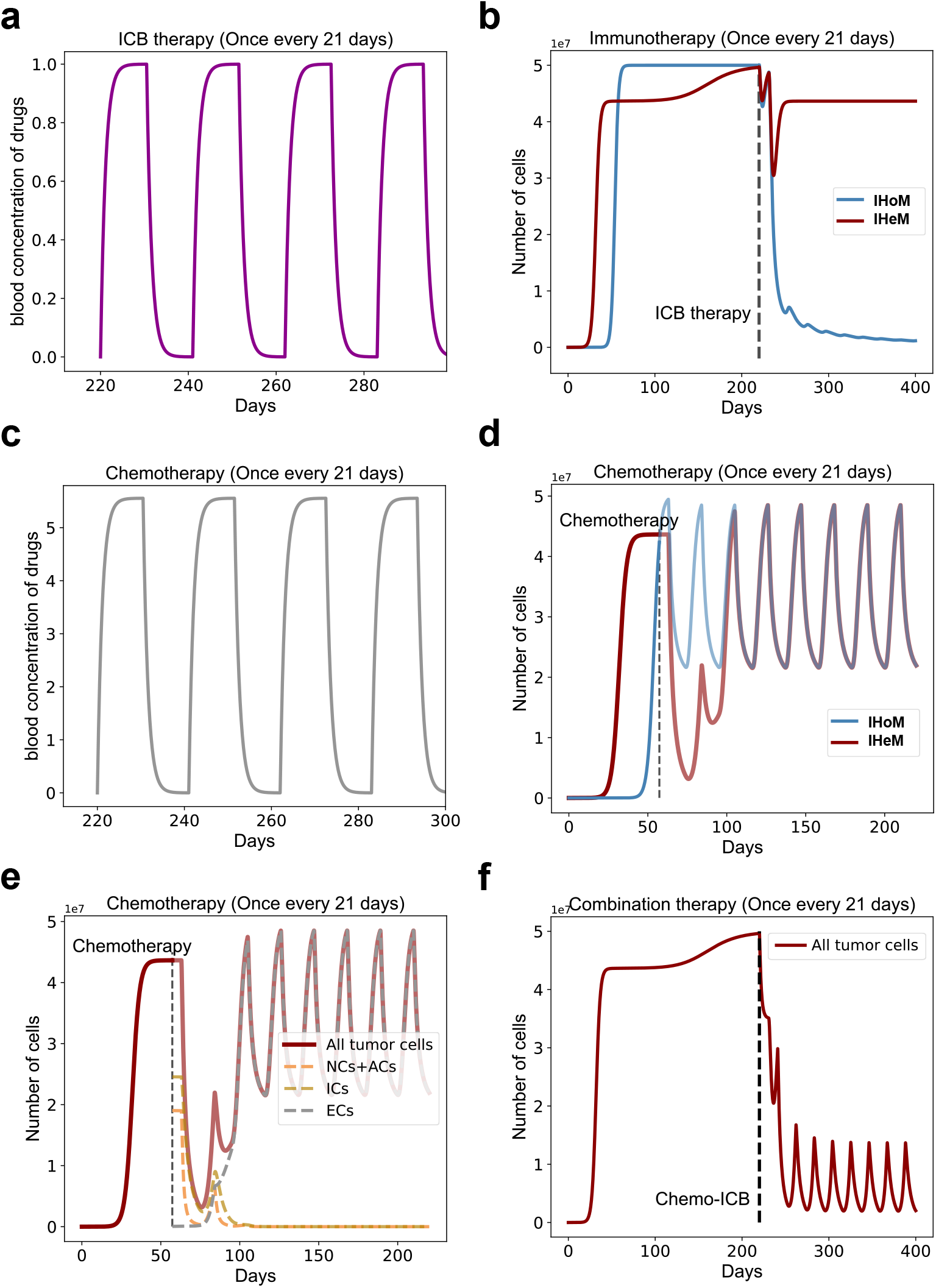
Responses to different therapies. **a**, Tumor growth curves under ICB therapy. **b**, Blood concentration change of ICB drug corresponding to treatment cycle. **c**, Tumor growth curves under chemotherapy. **d**, Blood concentration change of chemotherapy drug corresponding to treatment cycle. **e**, Growth curves of NCs+ACs, ICs and ECs under chemotherapy generated with numerical solutions. **f**, Tumor growth curves under combination therapy. Parameters: (*r*_1_, *r*_2_, *μ*_1_, *μ*_2_, *μ*_3_, *d, c, p*_*e*_, *f, γ, K*_*N*_, *K*_*A*_, *K*_*I*_, *K*_*E*_, *K*_*d*_, *v*_*m*_, *v*_*b*_) = (1, 0.5, 0.5, 0.6, 0.05, 0.5, 1 × 10^*−*8^, 1 × 10^*−*5^, 2*π/*21, 0.9, 0.9, 0.9, 0.9, 0.6, 0.6, 5, 0.9).

Initially, we examined the response of tumors under each model to single therapy. The results indicated that the tumor under the IHoM was responsive to ICB, leading to its extinction, while the tumor under the IHeM remained unresponsive, maintaining a high equilibrium population size (Fig. 5b). This outcome is consistent with previous findings [6–8]. In contrast to the IHoM, the tumor under the IHeM exhibited an early response followed by rapid relapse when treated with chemotherapy under the same strategy (Fig. 5d). To elucidate this phenomenon in the IHeM, we examined the dynamics of various cell components within the tumor. We observed a rapid increase in immune-escaped cells (ECs) after the initial response (Fig. 5e). The heightened sensitivity of non-cancerous cells (NCs) and antigenic cells (ACs) to chemotherapy led to their rapid decrease and extinction. Simultaneously, chemotherapy-induced damage to the immune system facilitated immune escape, providing ECs with a distinct survival advantage and resulting in rapid relapse.

Following the relapse, the tumor, initially evolving under the dynamics of the IHeM, now transitions to being governed by the IHoM. This transition results in a curative effect when combined with immune checkpoint blockade (ICB) therapy (Fig. 5f). This outcome aligns with previous findings suggesting that chemotherapy increases PD-L1 expression in cancers [46]. We have elucidated how chemotherapy can induce favorable immunological conditions in chemo-ICB combination therapy, emphasizing the significant responsiveness of the IHoM to ICB.

To validate the transformation of the IHeM to the IHoM during early remission under chemotherapy, we examined tumor antigenic heterogeneity before and after chemotherapy. Referring to Fig. 4b, we observed that remission under chemotherapy occurs before tumor immune escape. We simulated chemotherapy and recorded the neoantigen landscapes of the tumors. Cell antigenicity, defined as the sum of all neoantigens’ antigenicity for each cell, served as our metric. The simulation results demonstrated a decline in tumor antigenic Shannon diversity after chemotherapy (Fig. 6a). Additionally, the variations in tumor cell antigenicity distributions were significantly reduced post-chemotherapy (Fig. 6b and c). These findings suggest that tumors become more homogeneous after chemotherapy. Given that studies indicate antigenically homogeneous tumors are responsive to immune checkpoint blockade (ICB) therapy [6, 7, 47], our results unveil the underlying mechanism governing the response to ICB in combination with chemotherapy.

**Fig 6.**
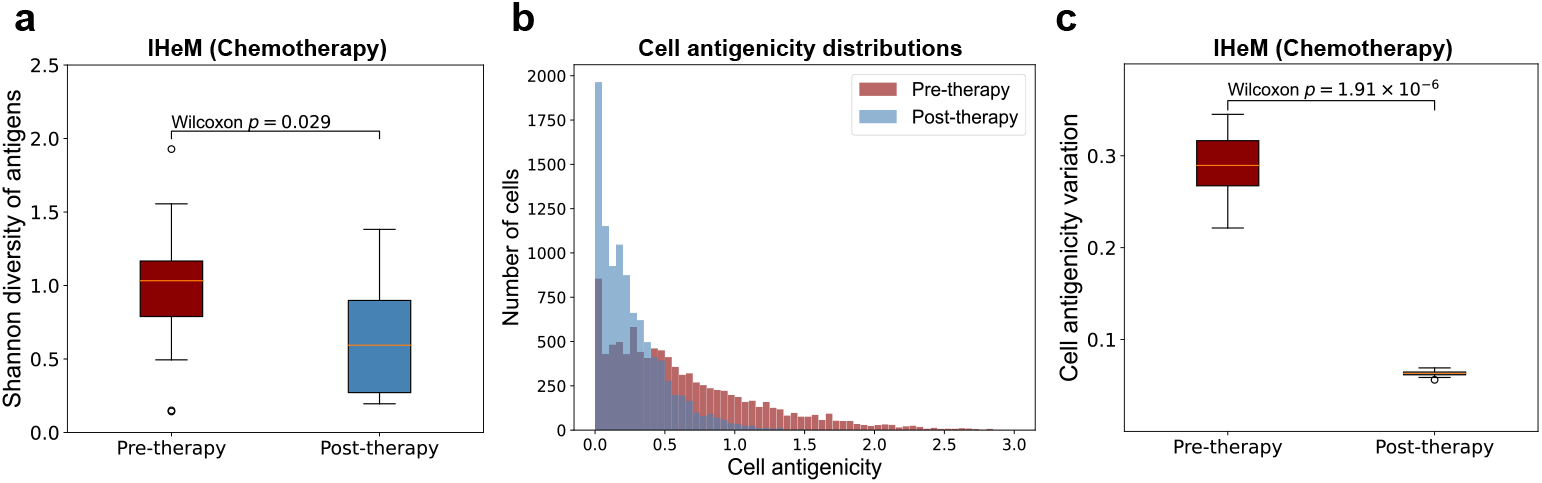
Effect of chemotherapy on antigens diversity. **a**, Box plots of Shannon diversities of neoantigens before and after 10 simulated chemotherapy. **b**, Distribution of cell antigenicity before and after one simulated chemotherapy. **c**, Box plots of cell antigenicity variation before and after 10 simulated chemotherapy.

The therapeutic response to drug combinations often improves with higher doses, but the consideration of patient tolerance during treatment is crucial. To explore the therapeutic effect of different dose pairs of drugs, we calculated the relative tumor size under each dose pair. Notably, a small dose of chemotherapy drug (*v*_*m*_ ≤ 1) combined with a certain ICB dose exceeding 0.5 demonstrated a significant therapeutic response (Fig. 7a). The use of a smaller chemotherapy dose causes less damage to the immune system while still modifying the tumor’s immunological conditions, thus showing promising effects in combination with ICB. This result underscores the pivotal role the immune system plays in combination therapy. Utilizing parameter values from Table 1, we conducted sensitivity analysis, revealing that our model results are not highly sensitive to changes in parameter values (Fig. 7b). This outcome validates the robustness of our conclusions and reinforces the reliability of our model’s predictions.

**Fig 7.**
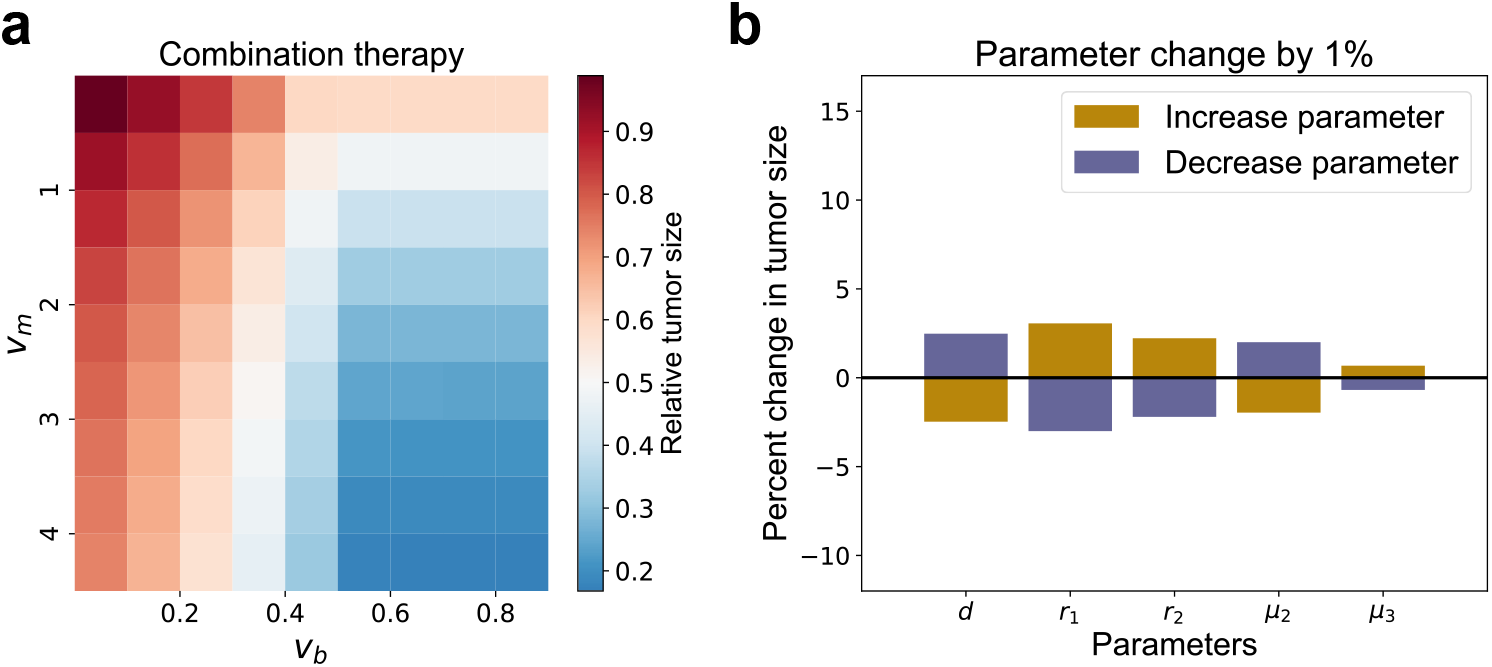
Effect of drug dose pair and sensitivity analysis. **a**, Therapeutic effect of different dose pairs of ICB and chemotherapy drugs. Therapeutic effect measured by relative tumor size (Post-therapy population size/Pre-therapy population size). **b**, Sensitivity analysis on parameters.

Based on the aforementioned results, the detrimental impact of chemotherapy on the immune system significantly influences the effectiveness of combination therapy. To mitigate the damage caused by chemotherapy to the immune system, we explored an alternative treatment protocol. In this revised approach, tumors were treated with chemotherapy during the first half of each treatment cycle, followed by immune checkpoint blockade (ICB) therapy in the second half (Fig. 8a). We assumed that this modified therapy protocol results in less immune system damage, introducing a healing factor *h*. Consequently, we adjusted the immune cell death rate from *d* · (1 − *K*_*I*_ (1 − *e*^*−M*^)) to *d* · (1 − *h K*_*I*_ (1 − *e*^*−M*^)), resulting in an improved combination therapy outcome (Fig. 8b). This restructured treatment approach leads to a mitigated relapse in each treatment cycle, ultimately culminating in the successful cure of cancer. Our findings suggest that even when the immune-damaging effects of chemotherapy may diminish the effectiveness of ICB treatment, maintaining the order of medication—chemotherapy preceding ICB treatment—remains a favorable strategy.

**Fig 8.**
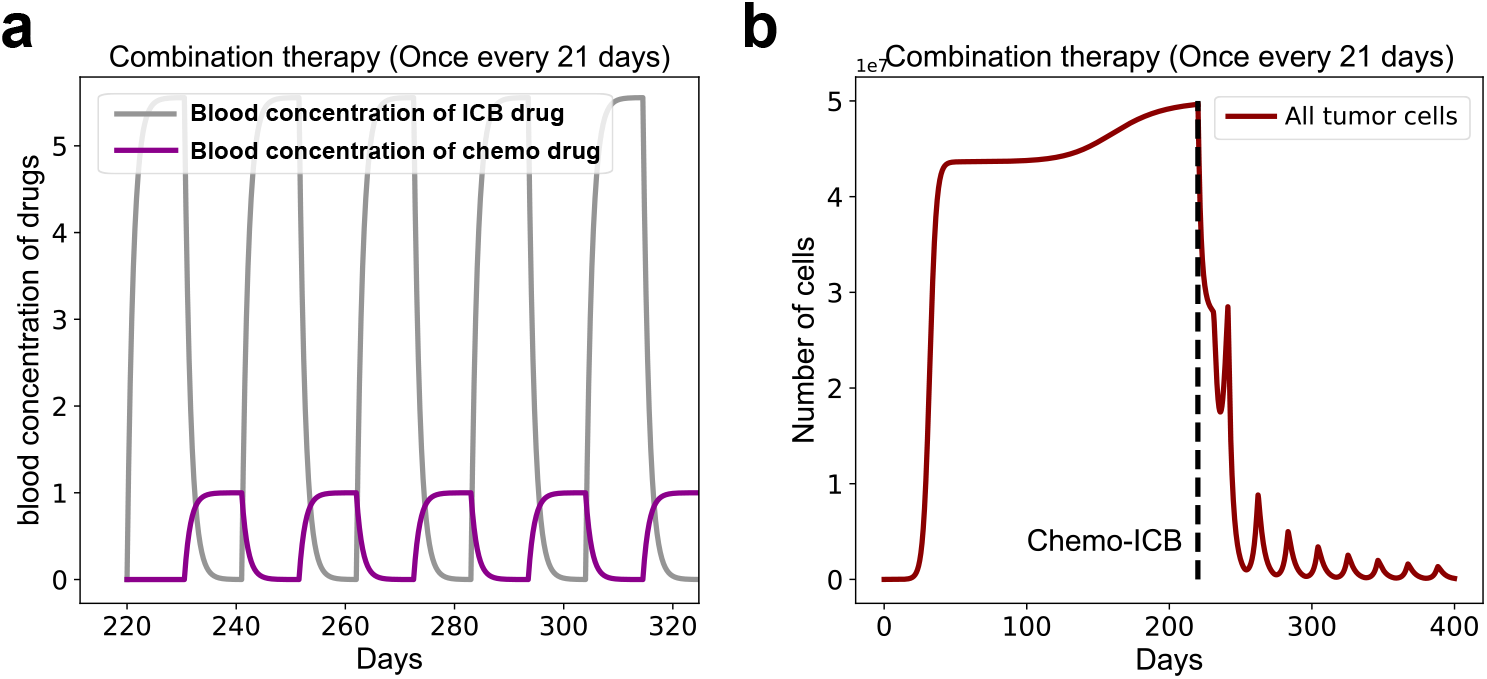
Revised combination therapy. **a**, Blood concentration curve of revised therapy protocol. **b**, Tumor growth curves under combination therapy under revised therapy protocol. Parameters: (*r*_1_, *r*_2_, *μ*_1_, *μ*_2_, *μ*_3_, *d, c, p*_*e*_, *f, γ, K*_*N*_, *K*_*A*_, *K*_*I*_, *K*_*E*_, *K*_*d*_, *v*_*m*_, *v*_*b*_, *h*) = (1, 0.5, 0.5, 0.6, 0.05, 0.5, 1 × 10^*−*8^, 1 × 10^*−*5^, 2*π/*21, 0.9, 0.9, 0.9, 0.9, 0.6, 0.6, 5, 0.9, 0.5).

## Discussion

In this study, we investigated the dynamics of cancer evolution during chemo-ICB therapy through the application of mathematical models. The adaptability of cancers to the immune system has been explored through various mechanisms [48–50]. By using our models, we illustrated an immune escape mechanism observed in antigenically heterogeneous tumors. This escape mechanism operates by regulating the proportions of antigenic cells (ACs) and immunogenic cells (ICs) in a frequency-dependent manner. The presence of such a mechanism aligns with reported instances of treatment failure in ICB, particularly in tumors characterized by a high burden of antigenic mutations that remain unresponsive to ICB [51]. Our assumptions, positing the origin of cancers from neutral cells (NC) with subsequent differentiation into ICs [35, 52–54], allowed us to explain how chemotherapy creates a conducive environment for tumor growth under immunogenic conditions. Despite the potential immune system damage caused by chemotherapy [33, 42, 43], an intriguing trade-off arises. This trade-off involves the transformation of the IHeM to the IHoM, ultimately rendering the tumor more amenable to ICB therapy. Our findings also highlight the enhanced therapy response observed in combination therapy scenarios. Furthermore, we conducted an analysis of prognostic factors such as cell proliferation rate and drug doses, shedding light on their influences within the context of the modeled therapeutic approaches.

The classical predator-prey model framework applied to cancer not only consider the growth dynamics of tumor cells but also account for the dynamics of effector cells in the tumor microenvironment, including T cells, NK cells, and Tregs cells, among others [33]. In our mathematical models, we quantify the impact of the tumor microenvironment using parameters, focusing on the intratumoral evolutionary mechanisms and characterizing the growth dynamics within the tumor-immune microenvironment. Moreover, our results reveal the significance of the tumor-immune microenvironment, particularly under the influence of chemotherapy. In terms of pharmacological toxicity, recent research has demonstrated that maintaining low concentrations of chemotherapeutic drugs and relatively high concentrations of immunotherapy drugs can effectively mitigate toxic effects [55], aligning with our own findings.

Our models considered the combination of chemotherapy and ICB therapy; however, the design of combination therapy typically involves selecting the optimal drug combination, determining the best dose pair, and establishing the order of drug delivery [24, 25, 56, 57]. While our models clarify general mechanisms underpinning therapies, we acknowledge that the intricacies of combination therapy design are multifaceted. We used parameters sourced from existing literature [4, 6, 16, 33, 34, 58, 59]. However, we encountered challenges in fitting our modeling results to actual patient data. Our models were constructed based on the assumption of cellular hierarchy and its applications to tumor immunology. Initially, our model assumed that antigenic cells (ACs) and immunogenic cells (ICs) follow distinct hierarchies of cell differentiation. However, real-world tumor scenarios may be more complex. We discussed the implications if the actual situation diverges from our hypothesis (Supplementary Fig. S1). Notably, as the proliferation rate of ACs diminishes relative to that of ICs, the differences in responses between ICB and chemotherapy decrease (Fig. S1a and b). Furthermore, responses to combination therapy decline as ACs become more differentiated than ICs (Fig. S1c). Despite these considerations, combination therapy still demonstrates an improved response compared to single treatments, underscoring its potential efficacy across various proliferation rate scenarios.

By uncovering general mechanisms of cancer evolution under therapeutic interventions, our models yield crucial insights for optimizing combination therapy strategies. The integration of chemotherapy and immunotherapy represents a promising avenue in cancer treatment, demonstrating positive outcomes in clinical trials. The next frontier involves refining chemo-ICB regimens based on synergistic mechanisms to enhance clinical efficacy further. Our models provide a framework for integrating drug combinations with patient data in future applications. Additionally, the optimal drug delivery order in combination alternate therapy is a pivotal consideration in treatment strategy optimization. Our work addresses this question, asserting that the order of medication administration should prioritize chemotherapy before ICB treatment. This finding contributes valuable guidance to the ongoing efforts in tailoring effective combination therapies for cancer patients.

## Supporting information

**Fig S1.**
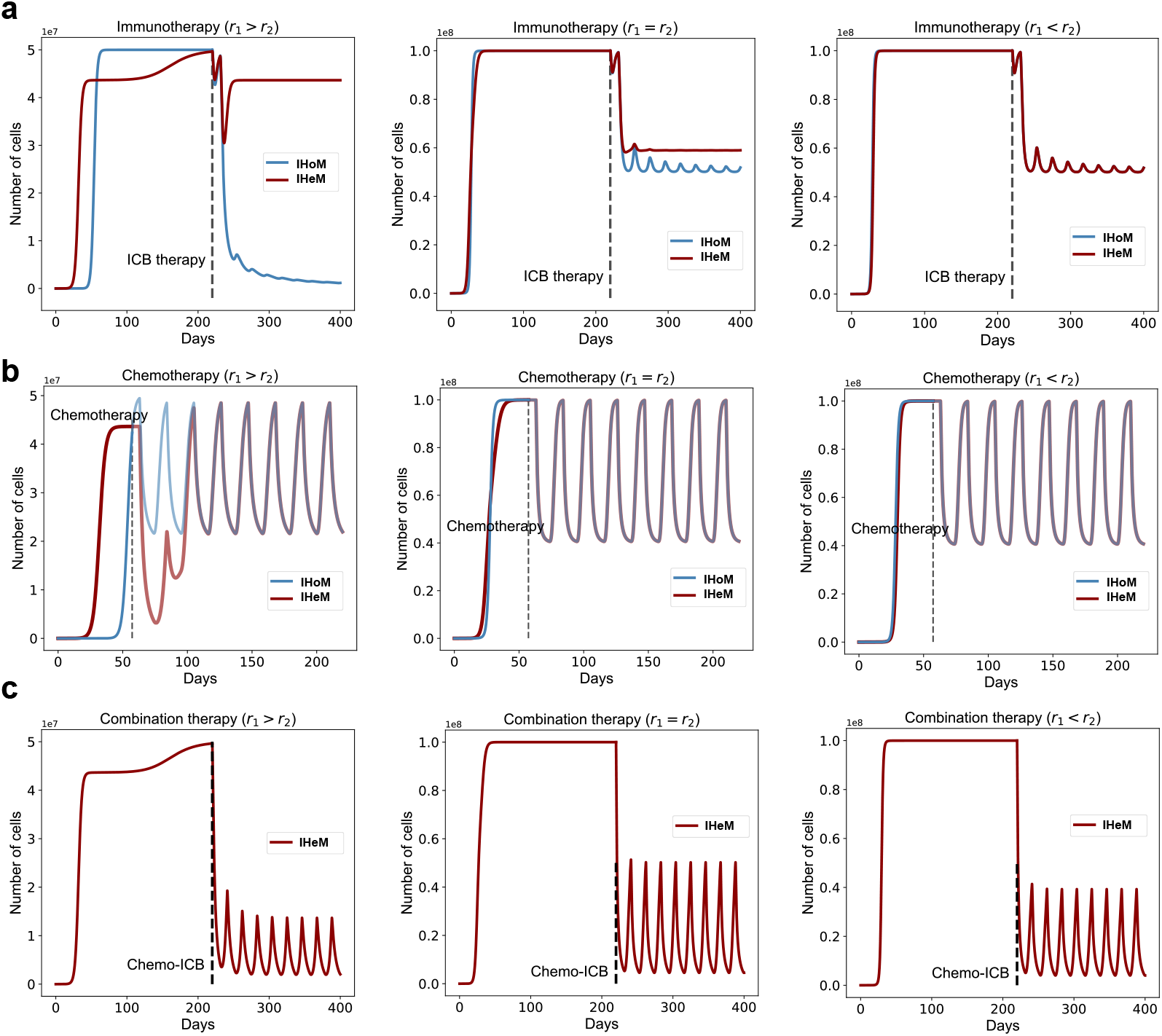
Responses to therapies in different scenarios. **a**, Tumor growth curves under ICB therapy. **b**, Tumor growth curves under chemotherapy. **c**, Tumor growth curves under combination therapy.Parameters: (*r*_1_, *r*_2_, *μ*_1_, *μ*_2_, *μ*_3_, *d, c, p*_*e*_, *f, γ, K*_*N*_, *K*_*A*_, *K*_*I*_, *K*_*E*_, *K*_*d*_, *v*_*m*_, *v*_*b*_) = (1, 0.5, 0.5, 0.6, 0.05, 0.5, 1 × 10^*−*8^, 1 × 10^*−*5^, 2*π/*21, 0.9, 0.9, 0.9, 0.9, 0.6, 0.6, 5, 0.9). *r*_1_ *> r*_2_ : (*r*_1_, *r*_2_) = (1, 0.5), *r*_1_ = *r*_2_ : (*r*_1_, *r*_2_) = (1, 1), *r*_1_ *< r*_2_ : (*r*_1_, *r*_2_) = (0.5, 1)

## Acknowledgments

We thank members of Zhou and Hu laboratories for constructive discussions. This work was supported by National Natural Science Foundation of China (11971405 to D.Z., 32270693 82241236 to Z.H.), Guangdong Basic and Applied Basic Research Foundation (2021B1515020042 to Z.H.) and Fundamental Research Funds for the Central Universities (20720230023 to D.Z.).

## Author Contributions

Conceptualization: Shaoqing Chen, Zheng Hu, Da Zhou

Formal analysis: Shaoqing Chen

Methodology: Shaoqing Chen, Zheng Hu, Da Zhou

Software: Shaoqing Chen

Supervision: Zheng Hu, Da Zhou

Visualization: Shaoqing Chen

Writing – original draft: Shaoqing Chen, Da Zhou

Writing – review editing: Shaoqing Chen, Zheng Hu, Da Zhou

## References

1. Coulie, Pierre G. et al. Tumour antigens recognized by T lymphocytes: at the core of cancer therapy. Nat. Rev. Cancer. 2014 Jan; 14(2):135–146.

2. Xiao, Yanping and Freeman Gordon J. The Microsatellite Instable Subset of Colorectal Cancer Is a Particularly Good Candidate for Checkpoint Blockade Immunotherapy. Cancer Discov. 2015 Jan; 5(1):16–18.

3. Luksza, Marta. et al. A neoantigen fitness model predicts tumour response to checkpoint blockade therapy. Nature. 2017 Nov; 551(7681):517–520.

4. Butner, Joseph D. et al. Early prediction of clinical response to checkpoint inhibitor therapy in human solid tumors through mathematical modeling. eLife. 2021 Nov; 10:e70130.

5. Auslander, Noam. et al. Robust prediction of response to immune checkpoint blockade therapy in metastatic melanoma. Nat. Med. 2018 Aug; 24(10):1545–1549.

6. Aguadé-Gorgorió, Guim and Solé, Ricard. Tumour neoantigen heterogeneity thresholds provide a time window for combination immunotherapy. J R Soc Interface. 2020 Oct; 17(171):20200736.

7. Wolf, Y. et al. UVB-induced tumor heterogeneity diminishes immune response in melanoma. Cell. 2019 Sep; 179(1):219–235.

8. Marusyk, Andriy and Almendro, Vanessa and Polyak, Kornelia. Intra-tumour heterogeneity: a looking glass for cancer?. Nat. Rev. Cancer. 2012 Apr; 12(5):323–334.

9. H. Schreiber and T.H. Wu and J. Nachman and W.M. Kast. Immunodominance and tumor escape. Semin. Cancer Biol. 2002 Feb; 12(1):25–31.

10. Gejman, Ron S. et al. Rejection of immunogenic tumor clones is limited by clonal fraction. eLife. 2018 Nov; 7:e41090. bibitemr2 Gejman, Ron S. et al. Rejection of immunogenic tumor clones is limited by clonal fraction. eLife. 2018 Nov; 7:e41090.

11. Hanahan, D., Weinberg, R. A. Hallmarks of cancer: the next generation. Cell. 2011. Fed; 144(5): 646–674.

12. Le, Dung T. et al. Mismatch repair deficiency predicts response of solid tumors to PD-1 blockade, Science. 2017. Jun; 357(6349): 409–413.

13. Alsaab H O, Sau S, Alzhrani R. et al. PD-1 and PD-L1 checkpoint signaling inhibition for cancer immunotherapy: mechanism, combinations, and clinical outcome. Front Pharmacol. 2017. Aug; 8: 561.

14. Mandal, Rajarsi. and others et al. Genetic diversity of tumors with mismatch repair deficiency influences anti-PD-1 therapy response. Science. 2019. May; 364(6439): 485–491.

15. Zapata, Luis. et al. Immune selection determines tumor antigenicity and influences response to checkpoint inhibitors. Nat. Genet. 2023 Mar; 55(3):451–460.

16. Mumenthaler, Shannon M. et al. Evolutionary modeling of combination treatment strategies to overcome resistance to tyrosine kinase inhibitors in non-small cell lung cancer. Mol. Pharm. 2011 Oct; 8(6): 2069–2079.

17. Kessler D A, Austin R H, Levine H. Resistance to chemotherapy: patient variability and cellular heterogeneity. Cancer research. 2014 Sep; 74(17): 4663–4670.

18. Heinhuis, KM and Ros, W and Kok, M and Steeghs, N and Beijnen, JH and Schellens, JHM. Enhancing antitumor response by combining immune checkpoint inhibitors with chemotherapy in solid tumors. Ann. Oncol. 2019 Feb; 30(2): 219–235.

19. Liu, Zhichao and Liu, Jun and Li, Zhigang. Chemoimmunotherapy in advanced esophageal squamous cell carcinoma: optimizing chemotherapy regimens for immunotherapy combination. Signal Transduction and Targeted Therapy. 2022 Jul; 7(1):233.

20. Dafni, Urania and Tsourti, Zoi and Vervita, Katerina and Peters, Solange. Immune checkpoint inhibitors, alone or in combination with chemotherapy, as first-line treatment for advanced non-small cell lung cancer. A systematic review and network meta-analysis. Lung cancer. 2019 Aug; 134: 127–140.

21. Machiels, Jean-Pascal H. et al. Cyclophosphamide, doxorubicin, and paclitaxel enhance the antitumor immune response of granulocyte/macrophage-colony stimulating factor-secreting whole-cell vaccines in HER-2/neu tolerized mice. Cancer Res. 2001 May; 61(9): 3689–3697.

22. Sakai, Hiromi. et al. Effects of anticancer agents on cell viability, proliferative activity and cytokine production of peripheral blood mononuclear cells. J Clin Biochem Nutr. 2012 Nov; 52(1): 64–71.

23. Pol, Jonathan. et al. Trial Watch: Immunogenic cell death inducers for anticancer chemotherapy. Oncoimmunology. 2015 Jan; 4(4):e1008866.

24. Lai, Xiulan. et al. Modeling combination therapy for breast cancer with BET and immune checkpoint inhibitors. Proc. Natl. Acad. Sci. U.S.A. 2018 May; 115(21): 5534–5539.

25. Lai, Xiulan and Friedman, Avner. Combination therapy of cancer with cancer vaccine and immune checkpoint inhibitors: A mathematical model. PLoS One. 2017 May; 12(5):e0178479.

26. Zhang, Haifeng and Lei, Jinzhi. Optimal treatment strategy of cancers with intratumor heterogeneity. Math Biosci Eng. 2022 Sep; 19(12): 13337–13373.

27. Ma, Shizhao and Lei, Jinzhi and Lai, Xiulan. Modeling tumour heterogeneity of PD-L1 expression in tumour progression and adaptive therapy. Math Biosci Eng. 2023 Jan; 86(3):38.

28. Deboever, Nathaniel and Zhang, Jianjun. Neoadjuvant chemo-immunotherapy for lung cancer: how much is too much?. Transl Lung Cancer Res. 2022 Jul; 11(12):2360.

29. Gong, Ke. et al. PLAC8 overexpression correlates with PD-L1 upregulation and acquired resistance to chemotherapies in gallbladder carcinoma. Biochem Bioph Res Co. 2019 Aug; 516(3): 983–990.

30. Jiao, Yuyan and Liu, Ming and Luo, Ningning and Guo, Hao and Li, Jianzhe. Successful treatment of advanced pulmonary sarcomatoid carcinoma with the PD-1 inhibitor toripalimab: a case report. Oral Oncol. 2021 Jan; 112: 104992.

31. Herbst R S. et al. Predictive correlates of response to the anti-PD-L1 antibody MPDL3280A in cancer patients. Nature. 2014 Nov; 515(7528): 563–567.

32. Gerard C L, Delyon J, Wicky A, et al. Turning tumors from cold to inflamed to improve immunotherapy response. Cancer Treat Rev. 2021 May; 101: 102227.

33. Eladdadi, Amina and Kim, Peter and Mallet, Dann. Mathematical models of tumor-immune system dynamics. Springer. 2014; 107.

34. Lakatos, E. et al. Evolutionary dynamics of neoantigens in growing tumors. Nat. Genet. 2020 Sep; 52(10):1057–1066.

35. Chen, Shaoqing and Xie, Duo and Wang, Jiguang and Hu, Zheng and Zhou, Da. Frequency-dependent selection of neoantigens fosters tumor immune escape and predicts immunotherapy response. bioRxiv. 2023 Aug.

36. Huang, Weini and Hauert, Christoph and Traulsen, Arne. Stochastic game dynamics under demographic fluctuations. Proc. Natl. Acad. Sci. 2015 Jul; 7112(29):9064–9069.

37. Baker, Nicholas E. Emerging mechanisms of cell competition. Nat. Rev. Genet. 2020 Aug; 21(11): 683–697.

38. Samoilenko, Anatolii Mikhailovich and Perestyuk, NA. Impulsive differential equations. World Scientific. 1995.

39. Eyer, Charles L. Goodman and Gilman’s the pharmacological basis of therapeutics. Am J Pharm Educ. 2002 66(1), 95.

40. Mitchison, Timothy J. The proliferation rate paradox in antimitotic chemotherapy. Mol. Biol. Cell. 2012 Oct; 23(1): 1–6.

41. Gardner, Shea N. A mechanistic, predictive model of dose-response curves for cell cycle phase-specific and-nonspecific drugs. Cancer Res. 2000 Mar; 60(5): 1417–1425.

42. Zhang, Yuanyuan. et al. Single-cell analyses reveal key immune cell subsets associated with response to PD-L1 blockade in triple-negative breast cancer. Cancer Cell. 2021 Dec; 39(12): 1578–1593.

43. Ghiringhelli, François and Apetoh, Lionel. The interplay between the immune system and chemotherapy: emerging methods for optimizing therapy. Expert Rev Clin Immunol. 2014 Dec; 10(1): 19–30.

44. Luster, M. I., Blank, J. A., and Dean, J. H. . Molecular and cellular basis of chemically induced immunotoxicity. Ann. Rev. Pharmacol. and Toxicol. 1987; 27(1): 23–49.

45. Principe, Nicola. et al. Comprehensive testing of chemotherapy and immune checkpoint blockade in preclinical cancer models identifies additive combinations. Front. Immunol. 2022 May; 13: 872295.

46. Ng, Hoi Yan. et al. Chemotherapeutic treatments increase PD-L1 expression in esophageal squamous cell carcinoma through EGFR/ERK activation. Transl. Oncol. 2018 Dec; 11(6): 1323–1333.

47. Reuben, Alexandre. et al. Genomic and immune heterogeneity are associated with differential responses to therapy in melanoma. NPJ Genom. Med. 2017 Apr; 2(1):10.

48. Gavin P. Dunn, Allen T. Bruce, Hiroaki Ikeda, Lloyd J. Old Robert D. Schreiber. Cancer immunoediting: from immunosurveillance to tumor escape. Nat. Immunol. 2002 Nov; 3(11):991–998.

49. Woo, Seng-Ryong. et al. Immune Inhibitory Molecules LAG-3 and PD-1 Synergistically Regulate T-cell Function to Promote Tumoral Immune Escape. Cancer Res. 2012 Feb; 72(4):917–927.

50. Beatty, Gregory L and Gladney Whitney L. Immune Escape Mechanisms as a Guide for Cancer Immunotherapy. Clin. Cancer Res. 2015 Feb; 21(4):687–692.

51. Gong, Xutong and Karchin, Rachel. Clustering by antigen-presenting genes reveals immune landscapes and predicts response to checkpoint immunotherapy. Sci. Rep. 2023 Jan; 13(1):950.

52. Bonnet, Dominique and Dick John E. Human acute myeloid leukemia is organized as a hierarchy that originates from a primitive hematopoietic cell. Nat. Med. 1997 Jul; 3(7): 730–737.

53. Perez-Losada, Jesus and Balmain, Allan. Stem-cell hierarchy in skin cancer. Nat. Rev. Cancer. 2003 Jun; 3(6): 434–443.

54. O’Brien Catherine Adell and Kreso, Antonija and Dick John E. Cancer stem cells in solid tumors: an overview. Semin Radiat Oncol. 2009 Apr; 19(2): 71–77.

55. Sanyi Tang, Shuo Li,Biao Tang, Xia Wang, Yanni Xiao and Robert A. Cheke. Hormetic and synergistic efects of cancer treatments revealed by modelling combinations of radio - or chemotherapy with immunotherapy . BMC Cancer. 2023 Oct; 123:1040.

56. H. Haeno and M. Gonen and M. Davis and J. Herman and C. Iacobuzio-Donahue and F. Michor. Computational Modeling of Pancreatic Cancer Reveals Kinetics of Metastasis Suggesting Optimum Treatment Strategies. Cell. 2012 Jan; 148(1): 362–375.

57. F. Michor and K. Beal. Improving Cancer Treatment via Mathematical Modeling: Surmounting the Challenges Is Worth the Effort. Cell. 2015 Nov; 163(5): 1059–1063.

58. Calabresi, Paul. Basic principles and clinical management of cancer. New York: McGraw-Hill. 1993

59. Perry, Michael Clinton. The chemotherapy source book. Lippincott Williams & Wilkins. 2008.

